# Characterizing the lipid-protein interface of the human serotonin transporter by crosslinking mass spectrometry

**DOI:** 10.1101/2020.06.01.128025

**Authors:** A.G. DeMarco, N.A. Ferraro, K. Sweigard, M. Cascio

**Author notes:** Corresponding Author: Michael Cascio, Mellon 331A, Department of Chemistry and Biochemistry, Duquense University, 600 Forbes Avenue, Pittsburgh, PA 15282.

## Abstract

Altered serotonin (5-HT) levels contribute to disease states such as depression and anxiety. Synaptic serotonin concentration is partially regulated by the serotonin transporter (SERT), making this transporter an important therapeutic target. This study seeks to examine the lipid accessible domains of hSERT to provide critical information regarding the apo-state of this transporter in a lipid environment. Recombinant hSERT was inducibly expressed in a human cell line. Solubilized SERT was purified by affinity chromatography using a FLAG Tag and reconstituted into mixed lipid vesicles containing our photoactivatable lipid probe. The lipid-accessible domains of the reconstituted transporter in membranes in its apo-state were probed via photocrosslinking to azi-cholesterol followed by quadrupole time of flight liquid chromatography-mass spectrometry (QTOF-LC-MS). MS studies identified crosslinks in three transmembrane loops consistent with the known topology of SERT. Surprisingly, the amino- and carboxy-terminal domains were similarly crosslinked by cholesterol, suggesting that these regions may also be intimately associated with the lipid bilayer. The data presented herein assist in further refining our understanding of the topography of the apo-state of hSERT via analysis of lipid accessibility.

## INTRODUCTION

Serotonin, dopamine, and norepinephrine are all monoamine neurotransmitters whose transporters are members of the neurotransmitter sodium symporter (NSS) family that includes the dopamine transporter (DAT), the serotonin transporter (SERT,) and the norepinephrine transporter (NET)(1). NSS uses electrochemical sodium and chloride gradients to drive neurotransmitter transport across the bilayer with the primary biological function of regulating monoamine transmission. Specifically, SERT governs a high-affinity uptake mechanism for serotonin (5-hydroxytryptamine, 5-HT) located on presynaptic 5-HT nerve terminals.

Given the role of NSS in recycling neurotransmitters and affecting their synaptic concentrations, NSS dysfunction is linked to neurological disorders and plays a significant role in their pathophysiology. SERT knockout mice demonstrate the role the protein plays in neuropathology (2,3). Further studies with knockout mice show epistasis between SERT and either DAT or NET (4), further enhancing the clinical complexity and making it difficult to assign specific roles of each NSS in diagnoses. Clinical manifestations of altered serotonin levels may be attributable to aberrant SERT function and may present as depression and anxiety (5). Current treatments typically include some permutation of monoamine transporter inhibitors such as selective serotonin reuptake inhibitors (SSRIs) or tricyclic antidepressants.

All NSS transporters possess 12 transmembrane helices with both the amino- and carboxy-termini (N-term and C-term, respectively) modeled in the cytoplasm. Historically, much of the structural information about the NSS family had been inferred from the known structure of LeuT (6-8) though there is only 20-25% amino acid homology between LeuT and members of NSS family (9,10). Despite this limited sequence homology, LeuT possesses a high structural homology in its agonist binding site when compared to NSS and has high affinity and is inhibited by SSRIs (8,11). The central binding site for leucine is approximately equidistant from both the extra- and intracellular face and contains both hydrophilic and hydrophobic amino acids. Since the determination of the LeuT structure, the structures of eukaryotic NSS family members in various liganded states have become available (12-15). These static structures provide essential information necessary to understand NSS activity at a molecular level, providing snapshots of various types of solubilized receptors in low energy states. However, an integral part of drug discovery is informed by a molecular understanding of protein allostery, the thermodynamic process in which proteins change conformation due to the interactions of chemically different secondary ligands (16,17). Allostery is a critical regulatory mechanism, controlling metabolic and signal transduction pathways.

Due to constraints of protein crystallography (18,19) it is oftentimes difficult to collate static crystallographic structures with the dynamics of molecular function. It is important to note that these structures are all of NSS transporters in detergent micelles and provide little direct data concerning specific lipid-protein interactions. In addition, many alterations were introduced in the sequences of the respective transporters for thermostability, limiting the functionality of the respective transporter and leaving sections of the transporter structurally unresolved. Allosteric modulation is not just restricted to soluble ligands, as studies of hDAT demonstrated that changes in cholesterol concentrations also drives conformational changes in the protein that ultimately affected ligand binding (20) and lipids can act in regulating transporter activity (21). To develop improved therapeutic modalities, therefore, studies refining the protein-lipid interface of hSERT in reconstituted vesicles in the absence of any ligands offer the possibility to contribute to our understanding of the structure of NSS transporters in their apo-state.

The lipid bilayer is a non-covalent assembly of proteins and lipids, and its spontaneous formation is driven by thermodynamic interactions between proteins and insoluble amphiphiles of which the bilayer is composed. The bilayer serves to minimize the entropic strain that arises when hydrophilic groups are forced to order themselves around hydrophobic structures. Most lipids in the bilayer are phospholipids, whose hydrophobic tails form a permeability barrier that protects the cell by compartmentalization and allowing for essential electrochemical gradients necessary for life processes. Lipid compositions have been shown to alter membrane protein function, either indirectly by affecting the physicochemical nature of the bilayer, or via specific lipid-protein interactions (22). hSERT studies examining lipid interactions have shown that phosphatidylinositol 4,5-bisphosphate lipids (23) and cholesterol (24) modulate transporter conformation and activity. *In vitro* studies in rat and monkey brains have also shown a broad effect of cholesterol levels on monoamine transporters (25).

An established method that directly identifies protein-lipid interactions is chemical crosslinking (XL). For example, cholesterol homologs were used to map sites of cholesterol interactions with the peripheral-type benzodiazepine receptor providing for the identification of putative consensus sites for cholesterol binding (26). Cholesterol photoaffinity probes have also characterized lipid-protein interactions with ion channels (27,28). The advent of mass spectrometry (MS) methodologies offers the opportunity to examine membrane protein structure (29) to more sensitively and accurately detect and map the interaction of lipids with membrane proteins (30-32). In this study, we conduct studies analogous to those used in examining cholesterol interactions with the glycine receptor (33) to directly probe the lipid accessible regions of hSERT in its apo-state. After photoactivation, quadrupole time of flight liquid chromatography-mass spectrometry (QTOF LC-MS) was used to identify tryptic fragments of the lipid-protein interface that were covalently modified by photoactivatable cholesterol. A significant number of these peptides were confirmed with secondary MS/MS analysis and will serve to refine the understanding of the lipid-protein interface in hSERT.

## MATERIALS AND METHODS

### Materials

Stably-transfected tetracycline-inducible human embryonic kidney (TREx-HEK293) cells that express hSERT with an N-terminal His and FLAG-tag with a Tobacco Etch Virus (TEV) protease site were provided by Dr. Hidehito Takayama (34). 7-Azi-cholesterol was purchased from American Radiolabeled Chemicals (Saint Louis MO). Egg and plant mixed phospholipid stocks were purchased from Avanti Lipids. Biobeads, 30% Acrylamide/Bis Solution 29:1 (3.3% C), 40% Acrylamide/Bis Solution 37.5:1, and goat anti-mouse antibody was purchased from Biorad (Hercules, CA). TEMED was purchased from JT Baker (Phillipsburg, NJ). Aprotinin (Apr) and 72% trichloroacetic acid (TCA) were purchased from MP Biomedicals LLC. TEV protease solution, 20x ProTEV buffer, ProTEV Protease, Fusion Protein were purchased from Promega. Nickel superflow resin was purchased from Qiagen (Hilden, GE. Dithiothreitol (DTT) was purchased from Rosche (Basel, CH). 50 and 15 mL falcon centrifuge tubes, benzamide, 0.1 M benzethonium chloride (BCl), FLAG® resin, anti-FLAG® peptide, mouse monoclonal ANTI-FLAG® M2 antibody, ethylene glycol-bis(2-aminoethylether)-N,N,N′,N′-tetraacetic acid (EGTA), and free cholesterol were purchased from Sigma Aldrich (St. Louis, MO). Modified Lowry protein assay reagent, 2 mg/mL bovine serum albumin (BSA), phenylmethyl sulfonyl fluoride (PMSF), digitonin 99% purity, acetonitrile (ACN), deoxycholate (DOC), ethylenediaminetetraacetic acid (EDTA), 2N Folin & Ciocalteu phenol reagent 1-butanol, Slide-A-Lyzer® Dialysis Cassette 3500 Molecular Weight Cut Off 3-12 mL capacity, Pierce Trypsin Protease MS Grade (90305), Dulbecco’s modified eagle medium (DMEM), penicillin-streptomycin (PS), L-glutamine, and blasticidine were purchased from Thermo-Fisher Scientific (Waltham, MA). Microcentrifuge tubes and cell culture flasks were purchased from VWR Scientific (Radnor, PA)

### TRex-HEK-293 hSERT Induction, Transfection, and Expression

The induction protocol was adapted from Takayama *et al*. (34). Briefly, cells were incubated under 5% CO_2_, 37°C in DMEM, 10% FBS, 2 mM L-glutamine, 1% PS and 5µg/mL blasticidine under 5% CO_2_, in a 37°C incubator. At approximately 60% confluence, the expression vector was stably transfected and incubated for 24 hours at 5% CO_2_, 37°C. hSERT expression was induced by addition of 1 µg/mL tetracycline, and after 48 hours, the cells were collected in phosphate-buffered saline (PBS) (10 mM PO_4_^3−,^ 137 mM NaCl, and 2.7 mM KCl, pH 7.4), homogenized for 15 strokes in a lysis buffer (10 mM TRIS-HCl, 1 mM EDTA, pH 7.4, and anti-proteolytic cocktail (APC: 1.6 µunits/mL aprotinin, 100 µM phenylmethanesulfonyl fluoride, 1 mM benzamidine, 100 µM benzethonium chloride). A typical preparation consisted of 4 plates with a concentration of at least 3 × 10^6^ cells/mL. Membranes were pelleted by centrifugation at 100,000xg for 20 minutes at 4°C. This process was repeated, and the membrane fraction was flash frozen and stored at -80°C.

### Solubilization and Purification of hSERT

Cell pellets were resuspended in 10 ml of lysis buffer containing 5 mM Tris-Cl, 5 mM EDTA, 5 mM EGTA, 10 mM DTT, and APC at pH 7.4 and sonicated (VWR Scientific Branson Sonifier 250 Microtip limit 7, duty cycle 60%) twice for 15 seconds on ice. Membranes were pelleted by ultracentrifugation at 300,000xg for 30 minutes at 4°C. The pellet was resuspended in 10 mL of lysis buffer with 300 mM NaCl. The sonication steps and centrifugation steps were repeated. Membranes were then solubilized using a solubilization buffer containing 5 mM EDTA, 5 mM EGTA, 1.0 mL (1.5 mg/mL) lipids, 25 mM pH 7.4 potassium phosphate buffer (KP_*i*_), 100 mM potassium chloride (KCl), 1% digitonin, 10 mM DTT, APC, and 1.0% deoxycholate and sonicated twice on ice (VWR Scientific Branson Sonifier 250 Microtip limit 7, Duty Cycle 60%) for 15 seconds, 6 times and then rocked at 4°C overnight. The sample was then centrifuged as previously described, and the supernatant was used in subsequent purifications. 1.0 mL of FLAG resin was equilibrated with 10 mL of Tris-buffered saline (TBS) (50 mM Tris-Cl, 150 mM NaCl, pH 7.4) by centrifugation for 1 minute at 4°C at 3200xg and repeated 3 times. Solubilized membranes were added to the resin and placed on a nutator for 2 hrs at 4°C to allow binding. The supernatant was removed, and the resin washed with excess TBS. A 1mL elution buffer (pH 7.4, wash buffer containing 0.25 mg/mL FLAG Tag peptide) added to the resin centrifuge tube and placed on a nutator at 4°C then centrifuged as before. The supernatant containing purified hSERT was stored at -20°C. To remove the FLAG affinity tag, aliquots were once again bound to the resin and eluted by targeted cleavage by incubation with TEV protease at 4°C overnight following manufacturer’s protocol. Protein concentration was determined using a modified Lowry assay (35) using BSA as a standard. Western blot analysis was also performed according to the procedure detailed in Takayama *et al*. (primary antibody: mouse anti-FLAG M2 -1:8000- and fluorescent coupled secondary antibody: goat anti-mouse-1:5000)(34).

### hSERT Reconstitution

Solubilized SERT samples were split into two equivalent fractions and liposomes containing 0.5 mL lipids (90% plant, 10% egg phospholipid stocks) and 2.4 mM cholesterol with and without 7.3 µM Azi-cholesterol (from concentrated 5 mg/mL stock in ethanol), respectively. Samples were sonicated (VWR Scientific Branson Sonifier 250 Microtip limit 7, Duty Cycle 60%) for 10 seconds twice on the ice and dialyzed in 3500 molecular weight cut off cassettes separately against excess 25 mM KP_*i*_ at pH 7.4, repeated twice over 5 days at 4°C. Additionally, the cholesterol/lipid solutions for the respective samples were added during the sonication steps of dialysis, wherein samples were sonicated again, and pelleted at 300,000xg for 35 min at 4°C. Upon completion, the reconstituted protein was stored in a 25 mM KP_*i*_ buffer at pH 7.4.

### Azi-Cholesterol Crosslinking and Trypsinolysis

Samples with or without added azi-cholesterol were subjected to UV light in quartz cuvettes on ice with an ultraviolet lamp 420 W Hg Arc lamp (Newport Model 97435-1000-1, 260-320 nm) at 6.3 amps, with 5 minutes of exposure and 5 minutes recovery repeated 5 times. Photo-activation was conducted on a bed of ice to minimize sample warming. 16 µl of respective samples were subjected to SDS PAGE (11% resolving, 5% stacking) to isolate the purified protein band and remove the unbound cholesterol and lipids. hSERT has an approximate mass of 70 kDa, and appropriate gel slices were excised. Gel slices were washed 3 times with 700 µL 50% methanol, 50% 50mM Am-BIC buffer (pH 7.4) and shaken for 45 minutes at 900 rpm (VWR Thermal Shake Touch). 750 µL of ACN was added to each tube for 30 minutes to dehydrate the gel slices. Samples were vacuum centrifuged (Eppendorf 5301Vacufuge Concentrator) until dry. 200 ng of trypsin in 500 µl of buffer was added to all samples on ice for 15 minutes, then shaken at 900 rpm overnight at 42°C. The supernatant was removed and stored. 300 µL of extraction buffer (0.1% formic acid in 50% ACN) was added to all samples and shaken for 35 minutes at 900 rpm, and this process was repeated twice, and the supernatants combined.

### MS and MSMS Analysis

MS methodology and peptide fragmentation were adapted from analogous cholesterol crosslinking studies conducted on the glycine receptor (33). Extracted peptides were resuspended in 50 µL of 50:50 ACN: H_2_O with 0.1% formic acid. Measurements were conducted using an Agilent 6530 ESI-QTOF-LC/MS with an Agilent HPLC-Chip II G2420-62006 ProtID-Chip-150 comprised of a 40 nL enrichment column and a 75 μm x 150 mm separation column packed with Zorbax 300SB-C18 5 μm material. The mass spectrometer was run in positive ion mode using internal standards (1221.9906 and 299.2944) for calibration. The nanoflow elution gradient was developed as follows at 0.50 µL/min of Solvent A (minute: percent A): 0.00: 95%; 3.00: 10%; 6.00: 70%; 9.00: 50%; 11.50: 95%; 13.00: 95%, with solvent A: 95% H_2_O; 5% ACN); and Solvent B: 5% H_2_O 95% ACN. Data were processed using Agilent Qualitative Analysis Software 6.0. Azi-cholesterol crosslinked peptides within a 10-ppm accuracy window were identified, allowing for oxidation and acrylamide modifications. Mass-shifted peptides containing crosslinked cholesterol were rerun under identical running conditions on the Agilent 6530 Q-TOF-MS, targeting the specific m/z ratio and retention time (RT) of the crosslinked peptides identified in initial MS analysis. Collision-induced dissociation (CID) was used for MS-MS fragmentation following a linear increase in collision energy by m/z using the equation: y=3.7x+2.5, where x is m/z and y, is the voltage. CID was performed at +/- 0.2 min from initial MS scan RT of each mass shifted precursor ion identified. Data were processed using Agilent Qualitative Analysis Software 6.0 in conjunction with ProteinProspector v5.14.2 available through the University of California, San Francisco.

## RESULTS

To examine, more fully, these interactions, cholesterol-hSERT interactions were studied using XL-MS. hSERT was over-expressed in a stably transfected HEK-293 cell line via tetracycline induction (34). One week after induction, the cell membranes were collected by centrifugation, and the hSERT in washed membranes was solubilized and purified by FLAG immunoprecipitation. After proteolytic removal of the N-terminal FLAG tag, eluted purified hSERT in mixed micelles containing mixed phospholipids and cholesterol at a concentration of ∼ 1.65 mg/ml, (with more than 10% of the lipid mass being cholesterol) was reconstituted into small unilamellar vesicles after detergent removal. In analogous studies using this overexpressing inducible cell line, purified hSERT retained nanomolar binding to SSRI analogs during purification (36) and Castellano *et al*. (manuscript in preparation). Experimental and control liposomes were spiked to contain ∼2.4 mM cholesterol with and without 7.3 µM azi-cholesterol, respectively, in addition to other mixed lipids. The average concentration of hSERT used in these studies was 267 +/- 1 µg/mL. Cholesterol has been shown to interact with membrane proteins and alter their function (37-41). as well as assist in the formation of lipid rafts (41,42). At > ∼35 mol%, cholesterol is considered to be chemically active in the membrane; below this level, cholesterol has negligible chemical activity and thus would be expected to interact non-specifically with proteins in the bilayer (43,44). The cholesterol interactions noted in this study occur at a cholesterol concentration of ∼15 mol%, where cholesterol is expected to have negligible chemical activity. At this reduced level of cholesterol, it is expected that the trace amounts of azi-cholesterol will probe the lipid accessible regions of the transporter in its apo-state and will act primarily as a stochastic probe for lipid accessibility (though even at its expected low chemical activity, we cannot rule out specific cholesterol binding sites are also crosslinked given the sensitivity of mass spectrometers). Typical differentiation of specific and non-specific binding sites as a function of ligand (in this case, cholesterol) concentration was not conducted as these studies would be confounded and be uninterpretable due to potential changes in hSERT conformation as a function of altered cholesterol levels, as well as changes in bilayer properties for example thickness, lateral pressure, and or fluidity; as a function of altered cholesterol composition.

Photoactivatable cholesterol analogs have been utilized to efficiently probe the lipid-accessible regions of transmembrane protein (27). The cholesterol analog used in this study includes a photoactivatable diarizine moiety at carbon seven (**Fig. 1**). Irradiation of this analog produces reactive carbene intermediates that form stable adducts with neighboring lipid or protein (37,38). Given the location of the carbene, it is expected that the cholesterol will label proteins at the top and bottom of the acyl chain region of the bilayer, close to the interfacial headgroup region.

**FIGURE 1.**
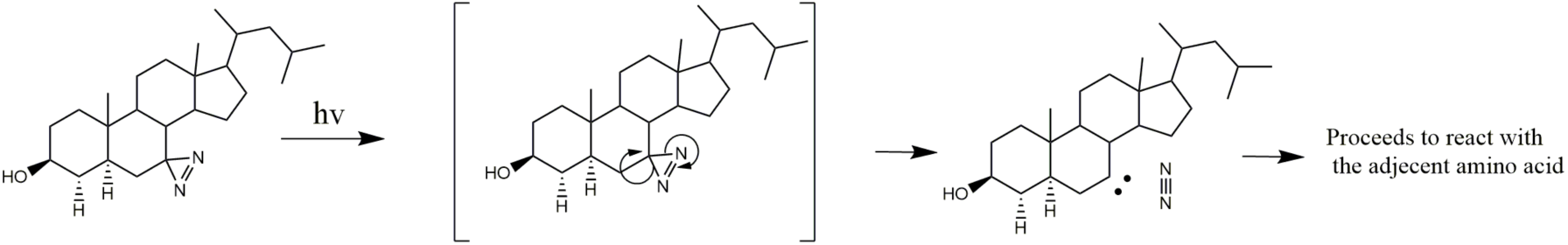
Azi-Cholesterol Photoactivation. Schematic cartoon showing the formation and location of the reactive carbene after UV irradiation.

After photoactivation, experimental and control samples (lacking azi-cholesterol) were subjected to SDS PAGE, and an in-gel trypsin digest of high MW bands corresponding to hSERT monomers and oligomers was performed. The extracted tryptic peptides were then analyzed by QTOF LC-MS, and only datasets with >20% coverage was selected for subsequent analyses by CID fragmentation. Overall, coverage of hSERT in these studies was ∼ 62% (see Supplementary Information for coverage map and non-compiled crosslinking Tables). Mass shifted tryptic mass ions after C18 LC chip separation were identified by MS. Cut-offs for precursor ion scans assignment used a mass error range of +/- 10 ppm. Crosslinking analyses was not limited to single cholesterol crosslinking event, and up to 4 cholesterol crosslinks per tryptic peptide were allowed, as cholesterol is distributed in both leaflets of the lipid bilayer and thus has the potential to interact at multiple sites within a single tryptic peptide, as well as potential “piggybacking” (cholesterol crosslinking to hSERT-bound cholesterol). From ∼20 picomoles of hSERT loaded on an SDS-PAGE and excised and trypsinized, 9 non-overlapping peptides were identified in 3 independent runs whose *m/z* were consistent with mass shifting by cholesterol crosslinking events (**Table 1**).

**TABLE 1:**
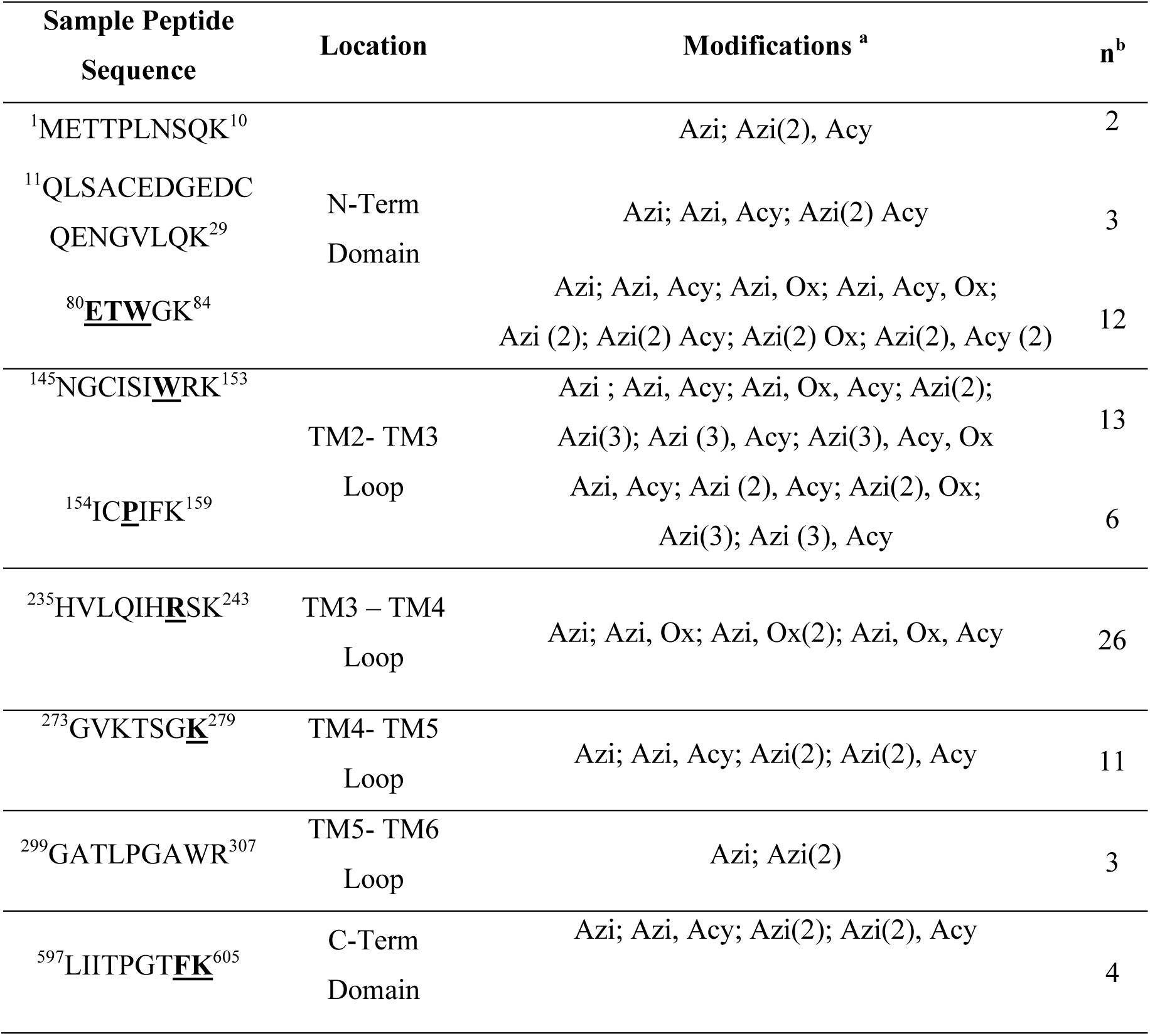
Cholesterol Crosslinking to hSERT. Compiled and condensed mass-shifted precursor ions crosslinked to cholesterol (within 10 ppm error) identified in at least 2 of 3 independent samples (each of which is provided in Supplementary Information). Single letter codes for the amino acids are listed along with their corresponding location in the hSERT structure. The bolded and underlined amino acids indicate the partial or complete resolution of the crosslinking site identified from CID product ions analyses. ^a^ Precursor ions modifications for each precursor ion are separated by semi-colons. Parenthesized numbers indicated the number of times that modification was observed in a given peptide. Acy is acrylamide, Ox is oxidation, Azi is covalent crosslinking from azi-cholesterol) ^b^Number of times the mass shifted peptide was identified across 3 independent samples

All mass shifted precursor ions containing crosslinked cholesterol were subsequently targeted for single fragmentation by CID in MS/MS studies, creating *a, b*, and *y* product ions that differed in mass depending on the site of bond breakage and charge localization (45). These product ion scans allow one to further refine the site of covalent cholesterol crosslinking within the tryptic peptide as mass-shifted product ions identify regions containing the attached cholesterol. Given that tryptic peptides could be modified at more than a single site, isobaric precursor ions can complicate and preclude assignment. To correct for this potential complication, MS/MS spectra are strictly matched by retention time to their precursor ion as in-line C18 liquid chromatography was used for potential separation of isobaric precursor mass ions. A 0.1 Da error cut-off was used in analyses of product ion scans. A representative full ion chromatograph, and a selected precursor and product ion scan, as well as the peptide assignment, is shown in **Fig. 2**. The site(s) of covalent cholesterol-binding could be refined within the peptide fragment by assignment of *a, b*, and *y* ions in product ion scans; however, the unambiguous assignment was not always possible in three cases. The assigned sites of covalent attachment are shown as bolded and underlined residues in **Table 1**.

**FIGURE 2:**
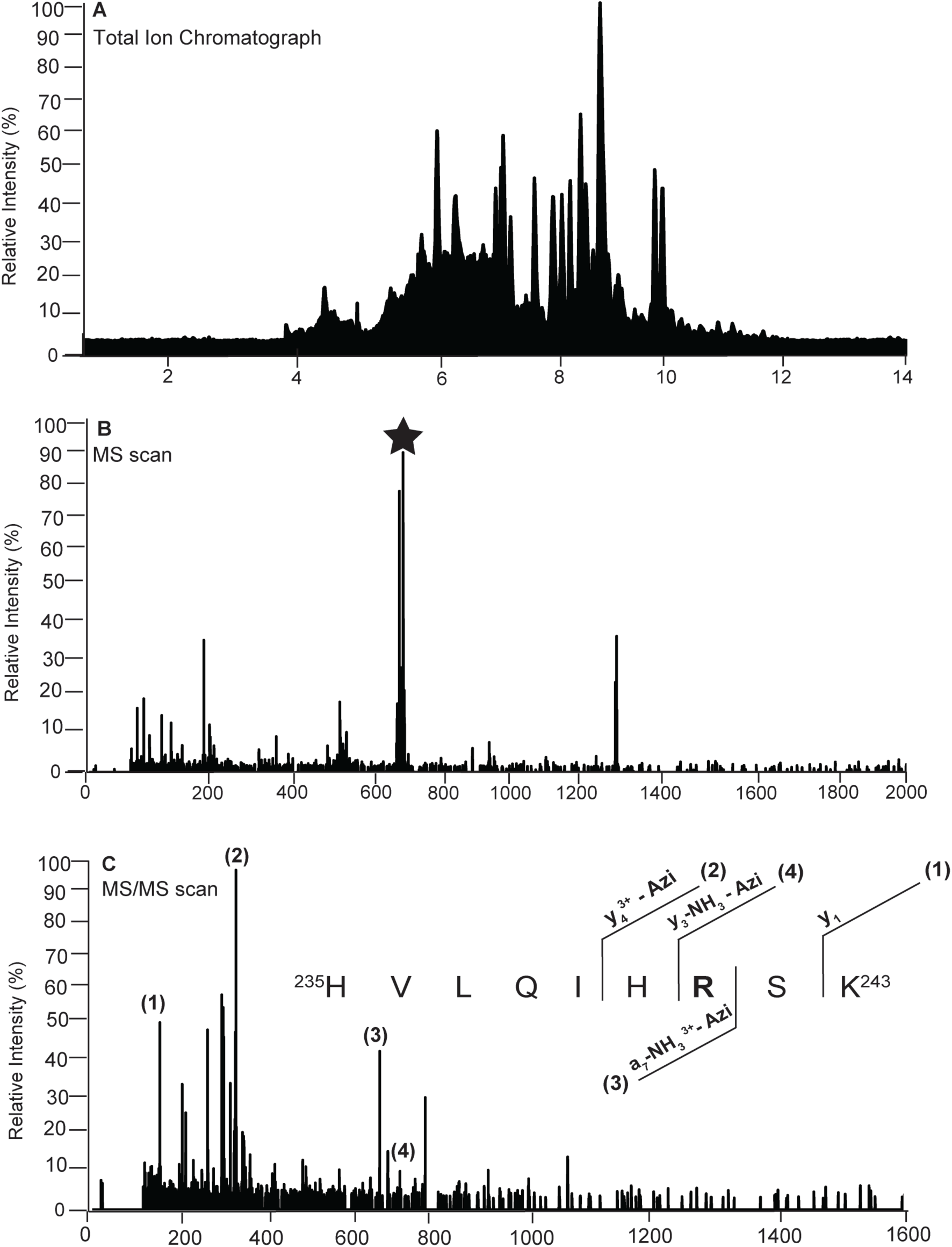
Representative MS and MS/MS spectrum of a single precursor and product ion scan. In a single trial run, from the full ion chromatograph (A), a precursor ion with an *m/z* 760.51 and retention time of ∼5.90 minutes (B) corresponding to a peptide ^235^HTVLQIH**R**SK^243^ mass shifted by a cholesterol adduct was subjected to CID fragmentation (C). Analysis of the product ions (1-4), including those mass shifted by cholesterol crosslinking (-Azi), are consistent with the site of cholesterol modification being limited to the arginine at position 241 (inset to panel C).

Sites of cholesterol crosslinking identified in targeted MS/MS refinement were mapped on a model of hSERT structure (**Fig. 3** and **4**). Of note, not all mass shifted peptides are shown in these figures as the crystal structure used as the basis of the model is a thermostable mutated form of hSERT with truncated N- and C-termini that are absent in the refined PDB structure. These regions are discussed at greater length in the *Discussion* and highlight the utility of CX-MS based approach as a complement to other biophysical methods in that it can provide state-dependent local structural information of regions of supramolecular complexes that are not available in other high-resolution structures and can be conducted in a native-like lipid bilayer in the absence of detergents.

**FIGURE 3.**
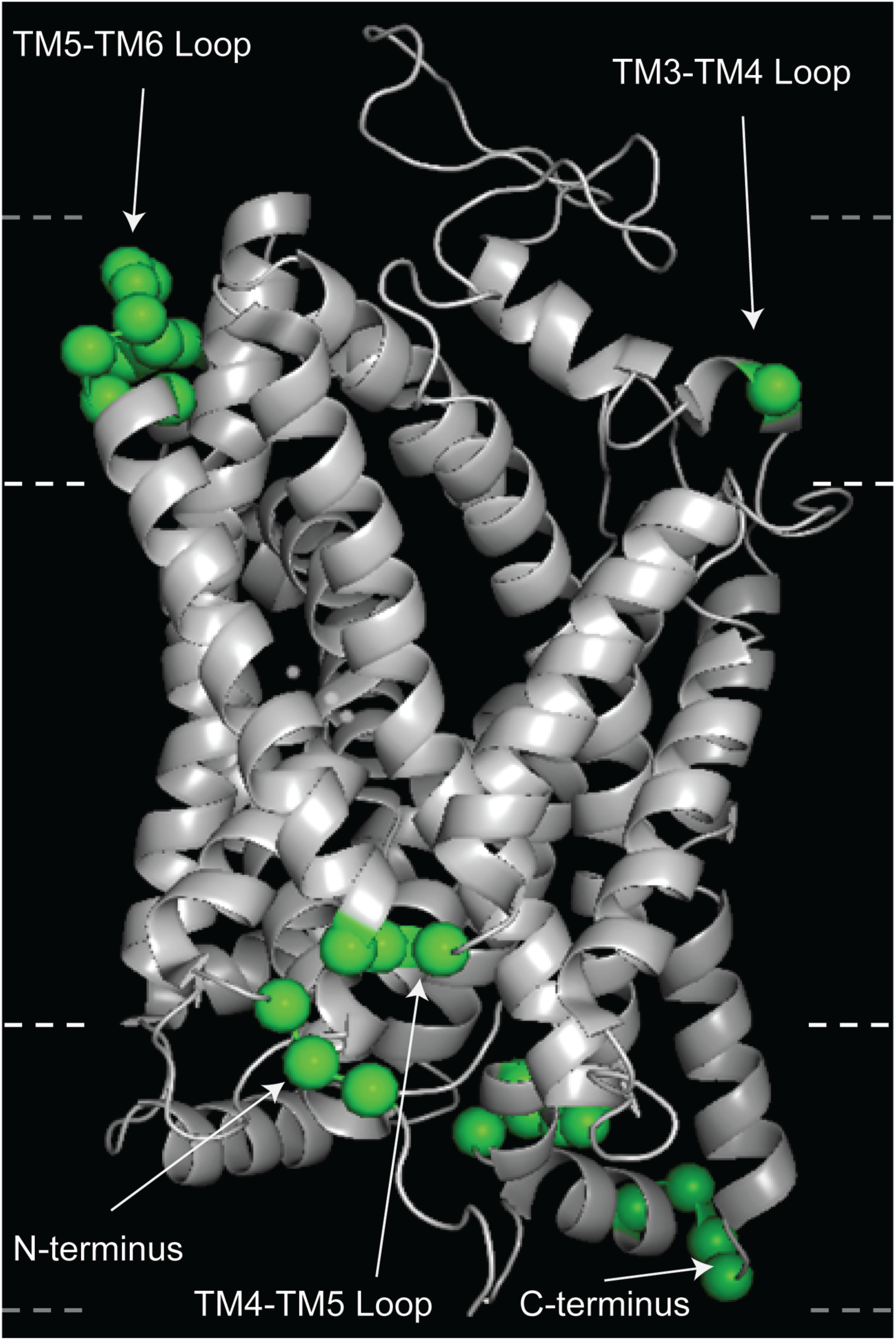
Sites of hSERT-cholesterol interactions as identified by MS-MS analyses. Refined sites of covalent crosslinking with azi-cholesterol (bolded and underlined amino acids in Table 1) mapped onto a schematic representation of hSERT (PDB #5I6X) with identified Cα atoms shown as green spheres. Dashed lines on edges of the image indicate approximate delimitations of the hydrophobic acyl chains (inner white dashed lines) and interfacial head group region (outer gray lines) of the bilayer and are provided for context. Figure was made using PyMol (57).

## DISCUSSION

Protein structure and/or oligomeric state can change as a function of lipid composition (23). For example, cardiolipin induces protein dimerization of LeuT (46), a paradigmatic monoamine transporter, and SERT structure and function is modulated by cholesterol (24). Thus, it is essential that complementary methodologies be developed to examine membrane protein structure in a lipid environment. The reconstituted bilayer composition was selected to represent a generic cell membrane containing a mix of phospholipids and cholesterol and does not represent the lipid bilayer composition of neuronal cells (which would have much higher concentrations of sphingomyelins). All of the cholesterol crosslinking studies reported in this study were conducted with cholesterol at ∼15 mol%, a concentration selected to allow identification of stochastic cholesterol interaction sites for the hSERT in its apo-state. At these concentration levels of cholesterol, cholesterol is expected to not have significant chemical activity and, therefore, our experimental conditions may preclude the detection of specific binding sites for cholesterol.

Coverage in both control and non-control studies is ∼62%, so it is expected that these studies do not exhaustively map all the lipid accessible regions of SERT as we cannot assume that all mass-shifted peptides may be detected. Regardless, these studies directly identify multiple direct cholesterol interactions localized to the N and C-termini, the TM2-TM3 and TM4-TM5 loops on the intracellular face, and the TM3-TM4 and TM5-TM6 loops on the extracellular face of hSERT. These interactions are highlighted in the schematic figure shown in **Fig. 3** (with the exception of crosslinking sites found at the N and C-termini, as these are absent in the model), but patterns of interactions may be similarly visualized by mapping these sites on a space-filling model of hSERT (**Fig. 4**). While these visual representations, based on the structure of hSERT with bound paroxetine (13), are expected to slightly differ from the apo structure of hSERT (which is currently not known) probed in our studies, the general structural details are expected to be fairly well conserved, and at the present time, the structural model used in mapping provides the best available template for visualization purposes. The pattern of crosslinking is strikingly restricted to the interfacial region of the membrane and the periphery of hSERT (**Figs. 3** and **4**). The depth of crosslinking is generally consistent with the expected location of the activated carbene in azi-cholesterol (**Fig. 1**) that is close to the hydroxyl headgroup of cholesterol and is similar in membrane depth to analogous studies conducted by our group in examining cholesterol interactions with the glycine receptor (33).

**FIGURE 4.**
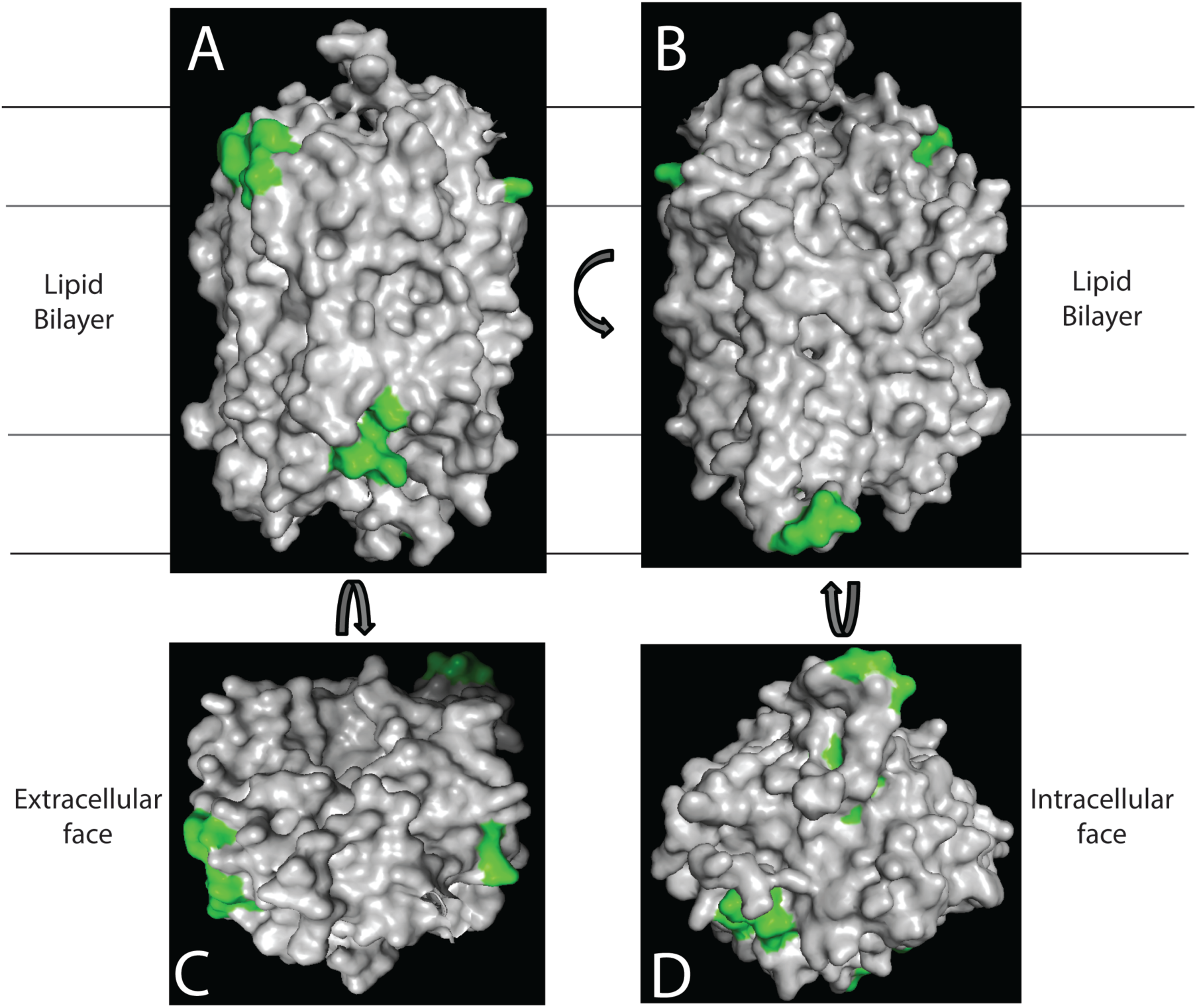
Sites of hSERT-cholesterol interactions mapped onto the space-filling model. Azi-cholesterol crosslinking sites (green) mapped on the space-filling model of hSERT (PDB #5I6X). **(A)** side-view of hSERT parallel with the membrane using the same orientation as in **Fig. 3. (B)** Side-view of hSERT rotated 180° from that shown in panel A. Inner horizontal lines indicate approximate region of hydrophobic interior of the bilayer (acyl chains) and outer lines indicate approximate limits for lipid head groups. **(C)** top-down (extracellular face) and **(D)** bottom-up (cytoplasmic face) views. Figure made using PyMol (57).

Of note, the loops connecting the transmembrane helices of MATs may not protrude far above the hydrophobic region of the bilayer and are probably involved in interactions with the polar head groups of the surrounding lipid and waters and ions that penetrate into the interfacial regions. Very little information is known about the structural and functional role of these loops, and their structure may be poorly represented in high-resolution studies that introduce thermostabilizing mutations and are conducted in non-physiological membrane-mimetic environments. Nevertheless, the labeling of the TM loops is consistent with other studies (discussed below), however crosslinking to the N and C-termini was unexpected as these regions are unresolved in current crystal structures and are portrayed in topological models as large intracellular loops (these loops, truncated in most structural studies, comprise about 20% of the molecular mass of this transporter).

It is interesting to compare the sites of cholesterol-SERT interactions observed in these studies with observed sites of interaction observed or predicted in other studies. In the crystal structure of hSERT bound to paroxetine in micelles, a cholesteryl ester was resolved in the crystal structure and was adjacent to TM12, in the extracellular leaflet (close to the TM11-12 loop). We did not observe cholesterol crosslinking interactions in this region but should not be considered as inconsistent with crosslinking-mass spectrometry (CX-MS) studies. As noted above, limited coverage in our CX-MS studies does not allow interpretation of negative results (e.g., the absence of observed crosslinks should not be interpreted as evidence of non-interactions), so we cannot ascertain if cholesterol similarly binds to this region in the apo-state. Consensus sequences for cholesterol binding have been identified (47-49). Cholesterol recognition amino acid consensus sequence (CRAC) is a linear motif (L/V)-X_1-5_-Y-X_1-5_-(K/R) where X is any hydrophobic amino acid, and the inversion of the CRAC domain referred to as the CARC, has consensus sequence of (K/R)-X_1-5_-(Y/F)-X_1-5_-(L/V). Both motifs are found in hSERT, and some of our identified sites of crosslinking overlap or are proximal to CARC and CRAC sequences. CRAC or CARC sites were also observed in the N-terminal domain (residues 85-92), the TM4-TM5 loop (residues 264-271), and TM5-TM6 loop (residues 309-314). A similar overlap between our experimental results and computational simulations of monoamine transporters (SERT, as well as dopamine and norepinephrine transporters) was observed by Zeppelin *et al*. (50), where it was speculated that cholesterol binding to the N-terminal domain/TM1a and TM5 (adjacent to the TM4TM5 loop) inhibits their necessary movement required for out-to-inward reverse transport. The interaction of cholesterol with TM1a and TM5 was subsequently confirmed by mutagenesis studies (24). The identification of crosslinking sites consistent with predicted and determined cholesterol-specific binding sites suggest that these studies are not strictly stochastic, and that even at low cholesterol availability used in our studies, CX-MS studies are sensitive enough to identify low-abundant binding events.

Our crosslinking data shows crosslinking of cholesterol to the region immediate proximal to TM1a (^80^**<u>ETW</u>**GK^84^), but also regions within peptides 1-10 and 11-29. In addition, crosslinking is also localized in MS/MS studies to residues 602-605 at the C–terminus. This suggests that these regions, unresolved in all published structures, are localized to the bilayer. Three other unique cholesterol interaction sites were identified in these studies in the TM2 – TM3 loop and the TM3-TM4 loop, more specifically, ^145^NGCISI**<u>W</u>**RK^153, 154^IC**<u>P</u>**IFK^159^, and ^235^HVLQIH**<u>R</u>**SK^241^. The latter is significant as the large extracellular TM3-TM4 loop is typically truncated when expressed for high-resolution studies, so very little information is known about the structure of this loop. Also, it is interesting to speculate on the potential role of membrane association, as the terminal loops of monoamine transporters are speculated to interact with phosphatidylinositol 4,5 bisphosphate (PIP_2_) and play a role in transporter regulation (51,52). These protein-lipid interactions interaction between SERT and PIP2 has been shown to be a prerequisite for amphetamine modulation of SERT function (53) and the N-terminus has been shown to be required for reverse transport mediated by amphetamines (54). Given the role of PIP_2_ binding and the presence of more negatively charged phospholipids in the inner leaflet, it is interesting to note that the crosslinked peptides presumed to be constrained to the inner leaflet contain a preponderance of basic amino acids (N- and C-termini, TM3-TM4 loop). These same regions contain sites of phosphorylation (55), raising the possibility that part of the functional effects of phosphorylation may be attributable to electrostatics-driven protein remodeling in the interfacial regions of the bilayer as a function of the phosphorylation state. Analogous protein-lipid electrostatic interactions have been ascribed to the remodeling of the ephrin receptors (56).

In summary, these studies directly map the lipid-accessible surface of SERT as the covalent crosslinking from azi-cholesterol is presumed to be non-specific and provide evidence of close association of the protein with its surrounding bilayer. These studies suggest novel interactions with the N- and C-termini, as well as TM2-TM3, TM3–TM4, TM4-TM5, and TM5-TM6 loops SERT in its apo-state. These studies also show the capability of CX-MS to map cholesterol-SERT interactions sensitively and accurately and foreshadow future studies wherein state-dependent crosslinking might be used to investigate the structure of the transporter in different allosteric states.

## AUTHOR CONTRIBUTIONS

A.G.D. and M.C. designed the research, A.G.D conducted the experiments, and N.A.F. assisted in MS experimental design and analyses. K.W. did visualization studies. A.G.D. and M.C. contributed to writing the paper, and N.A.F. provided editorial assistance.

## ACKNOWLEDGMENTS

The authors thank Kayce Tomcho, Emily Cooper, and Elizabeth Castellano for their technical support. This work was partially supported by funding from the National Science Foundation (NSF) Research Experiences for Undergraduate grant CHE-1659823. The purchase of the mass spectrometer was supported by the NSF (MRIDBI-0821401).

