## Supplemental Information for "Characterizing the lipid-protein interface of the human serotonin transporter by crosslinking mass spectrometry"

Independent Trial 1

| Peptide sequence | Location | Precursor Ion m/z | Precursor Ion retention time (min) | MS/MS ion matches |
|---|---|---|---|---|
| $^{80}$**ETW**GK$^{84}$ | | 1179.68 | 5.9 | $b_3$-$H_2O$ |
| $^{80}$ET**WGK**$^{84}$ | N-Term Domain | 1567.06 | 6.1 | $y^3$ |
| $^{80}$ETWG**K**$^{84}$ | | 1022.62 | 7.0 | $y_1^{2+}$, $y_2$-$NH_3$,$y^3$-$NH_3^{2+}$ |
| $^{145}$NGCISI**WR**$^{152}$ | | 1101.16 | 5.4 | $y_2^{2+}$ |
| $^{145}$NGCISIW**RK**$^{153}$ | | 1693.97 | 5.3 | $y_8$-$H_2O^{3+}$, $y_7$-$H_2O^{3+}$, $y_6^{2+}$, $y_3$-$NH_3$ |
| $^{145}$**NG**CISIWRKICPIF**K**$^{159}$ | TM2-TM3 Loop | 541.82 | 5.4 | $b_2$-$NH_3$, $y_9^{3+}$, $y_4$-$NH_3^{2+}$,$y_2^{2+}$, $y_1^{2+}$,$y_3^{2+}$ |
| $^{145}$NGCISIW**R**$^{152}$ | | 1338.81 | 6.0 | $y_2^{2+}$, $a_7$-$NH_3$ |
| $^{154}$IC**P**IFK$^{159}$ | | 1279.81 | 6.2 | $a_3$, .$a_2$, $y_2^{2+}$ |
| $^{235}$H**V**LQIHRS**K**$^{243}$ | TM3-TM4 Loop | 1694.07 | 7.3 | $y_8^{4+}$, $y_7^{2+}$, $y_6$-$H_2O^{3+}$, $y_5$-$H_2O$, $y_4$, $y_3$-$NH_3$, $y_1$, $a_8^{2+}$, $b_8$, $a_7^{3+}$, $b_6$, $a_5$,$b_5^{2+}$, $y_8^{4+}$ |
| $^{235}$**HVLQ**IHRSK$^{243}$ | | 1534.99 | 5.8 | $a_4$-$NH_3^{2+}$ |
| $^{275}$T**S**GK$^{279}$ | TM4-TM5 Loop | 952.66 | 5.7 | $y_2$-$NH_3$, $Y_3$-$NH_3^{2+}$, $b_2$-$H_2O$,$a_3$ |
| $^{597}$LIITPGT**FK**$^{605}$ | C-Term Domain | 458.31 | 5.5 | $y_2^{2+}$, $y_3^{2+}$, $y_4^{2+}$, $y_5^{2+}$, $y_6^{2+}$, $y_7^{2+}$ |

**Supplementary Table 1**: MS/MS ions table, where blue indicates a non-mass shifted ion. Bold and underlined residues indicate site/region of cholesterol attachment.

Independent Trial 2

| Peptide sequence | Location | Precursor Ion m/z | Precursor Ion retention time (min) | MS/MS ion matches |
|---|---|---|---|---|
| $^{80}$ET**W**GK$^{84}$ | N-Term Domain | 1006.65 | 6.0 | $y_1$, $y_3$-$NH_3^{2+}$, ,$a_2$,$b_2$-$H_2O$,$b_3$-$H_2O$ |
| $^{145}$NGCISI**WR**K$^{153}$ | TM2-TM3 Loop | 1012.17 | 5.4 | $y_8$-$NH_3^{2+}$,$y_6$-H2O$^{3+}$, $y_4$-$NH_3^{3+}$, $y_3$-$NH_3^{3+}$,$y_1^{2+}$,$b_8$ |
| $^{154}$I**CPIF**K$^{149}$ | | 1280.82 | 6.2 | $y_2$, $y_5$-$NH_3^{2+}$,$y_1$,$y_2$ |
| $^{235}$**HV**L**QI**HR$^{241}$ | TM3-TM4 Loop | 1304.87 | 5.5 | $y_5$-$NH_3^{2+}$, $y_4$-$NH_3^{2+}$, $a_2^{2+}$,$b_5$-$NH_3$, $b_6$ |
| $^{273}$GVKTSG**K**$^{279}$ | TM4-TM5 Loop | 541.72 | 6.4 | $y_1$-$NH_3^{3+}$,$b_5$-$H_2O^{2+}$, $a_4$-$NH_3^{2+}$,$b_6$-$NH_3^{2+}$ |
| $^{597}$LIIT**PGT**FK$^{605}$ | | 459.32 | 5.5 | $a_2$, $b_4$, $y_5$-$H_2O^{2+}$,$y_2$,$y_1$ |
| $^{597}$LIITPGTFKER**II**K$^{610}$ | C-Term Domain | 801.23 | 7.0 | $Y_{13}^{3+}$,$Y_{12}$-$NH_3^{4+}$,$y_{10}^{4+}$, $y_8$-$NH_3^{3+}$,$y_7$-$NH_3^{3+}$, $y_6$-$NH_3^{3+}$,$y_3$-$NH_3^{3+}$,$a_{13}^{3+}$ $a_{13}$-$NH_3^{3+}$ |

**Supplementary Table 2**: MS/MS ions table, where blue indicates a non-mass shifted ion. Bold and underlined residues indicate site/region of cholesterol attachment.

Independent Trial 3

| Peptide sequence | Location | Precursor Ion m/z | Precursor Ion retention time (min) | MS/MS ion matches |
|---|---|---|---|---|
| $^{145}$NGCISIW**R**K$^{153}$ | TM2-TM3 Loop | 1118.30 | 6.8 | $y_2$-$NH_3^{3+}$,$y^1$ |
| $^{145}$**N**GCISIWRK$^{153}$ | | 1118.31 | 7.3 | $MH^{2+,}$ $y_8$-$H_2O^{3+}$ ,$y_6^{2+}$,$y_3$ |
| $^{235}$H**V**LQI**HR**$^{242}$ | TM3-TM4 Loop | 1288.83 | 5.9 | $y_6$-$NH_3$, $y_2$-$NH_3$, $b_2^{2+}$,$a_2$,$b_5$+$H_2O$ |
| $^{235}$H**V**LQI**HR**$^{242}$ | | 1288.94 | 6.0 | $y_6$-$NH_3$, $y_2$,$a_2$,$b_5$ |
| $^{235}$**HVLQI**HRSK$^{243}$ | | 502.01 | 5.8 | $a_5^{2+}$, $b_5^{2+}$ |
| $^{235}$**HV**LQIH**R**SK$^{243}$ | | 768.51 | 5.8 | $a_2^{2+}$,$a_5^{2+}$,$a_7^{2+\cdot}$, $a_7$-$NH_3$, $y_3$-$NH_3$,$y_2$ |
| $^{235}$HVLQIH**R**SK$^{243}$ | | 760.51 | 5.9 | $y_4^{3+}$, $y_3$-$NH_3$,$y_1$, $a_7$-$NH_3^{3+}$ |
| $^{235}$H**V**LQIHR**S**K$^{243}$ | | 1549.00 | 5.9 | $y_8$-$NH_3^{3+}$, $y_7$-$H_2O^{2+}$, $y_7^{2+}$,$y_2$,$b_2$,$a_8$ |

**Supplementary Table 3**: MS/MS ions table, where blue indicates a non-mass shifted ion. Bold and underlined residues indicate site/region of cholesterol attachment.

Supplement Figure 1:

**METTPLNSQKQLSACEDGEDCQENGVLQKVVPTPGDKVESGQISNGYSAVPSPGAGDDTR**
**HSIPATTTTLVAELHQGERETWGKKVDFLLSVIGYAVDLGNVWR**FPYICYQNGGGAFLLPYT
IMAIFGGIPLFYMELALGQYHR**NGCISIWRKICPIFKGIGYAICIIAFYIASYYNTIMAWALYYLI**
**SSFTDQLPWTSCKNSWNTGNCTNYFSEDNITWTLHSTSPAEEFYTRHVLQIHRSKGLQDLG**
**GISWQLALCIMLIFTVIYFSIWKGVKTSGKVVWVTATFPYIILSVLLVRGATLPGAWRGVLF**
**YLKPNWQKLLETGVWIDAAAQIFFSLGPGFGVLLAFASYNKFNNNCYQDALVTSVVNCMT**
**SFVSGFVIFTVLGYMAEMRNEDVSEVAK**DAGPSLLFITYAEAIANMPASTFFAIIFFLMLITLGL
DSTFAGLEGVITAVLDEFPHVWAKRR**ERFVLAVVITCFFGSLVTLTFGGAYVVK**LLEEYATGP
AVLTVALIEAVAVSWFYGITQFCRDVKEMLGFSPGWFWRICWVAISPLFLLFIICSFLMSPPQLRL
FQYNYPYWSIILGYCIGTSSFICIPTYIAYR**LIITPGTFKERIIKSITPETPTEIPCGDIRLNAV**

Supplemental figure 1: Sequence coverage (62.1%) for all CX-MS trials. Bold amino acids indicate peptides precursor ions detected in analysis. Red amino acids indicate azi cholesterol modified sequences detected in 2 of 3 biological replicates.
